# Inflammaging mediates testosterone declines in men while maintaining high testosterone increases mortality risk

**DOI:** 10.64898/2026.06.04.730222

**Authors:** Jacob E. Aronoff, Benjamin C. Trumble

## Abstract

Later life is accompanied by testosterone declines alongside the development of chronic inflammation, termed inflammaging. A new theoretical model posits these processes are related through an energetic trade-off. As somatic damage accumulates, this should chronically activate the energetically costly inflammatory response. As an energy conserving response to promote cellular repair, testosterone production is expected to be suppressed. Consistent with this model, we find that markers of inflammaging, including IL-6 and GDF-15, mediate age-related testosterone declines in a large sample of male participants from the UK Biobank (n = 18,347, mean age 57 years, range 40-70). GDF-15, a marker of chronic inflammation and a key metabolic stress signaling protein, was the strongest predictor and mediator of testosterone declines. Further, individuals with high testosterone given their health and level of inflammation showed elevated mortality risk over follow up, consistent with a trade-off between maintenance and reproduction. Our results highlight the importance of considering energetic trade-offs to understand later life testosterone declines. They also highlight the importance of alleviating cellular damage that augments inflammaging and its down-stream hormonal effects. Finally, our study raises concern for exogenous testosterone therapies in the context of chronic inflammation, which could increase mortality risk.

**Graphical Abstract:** 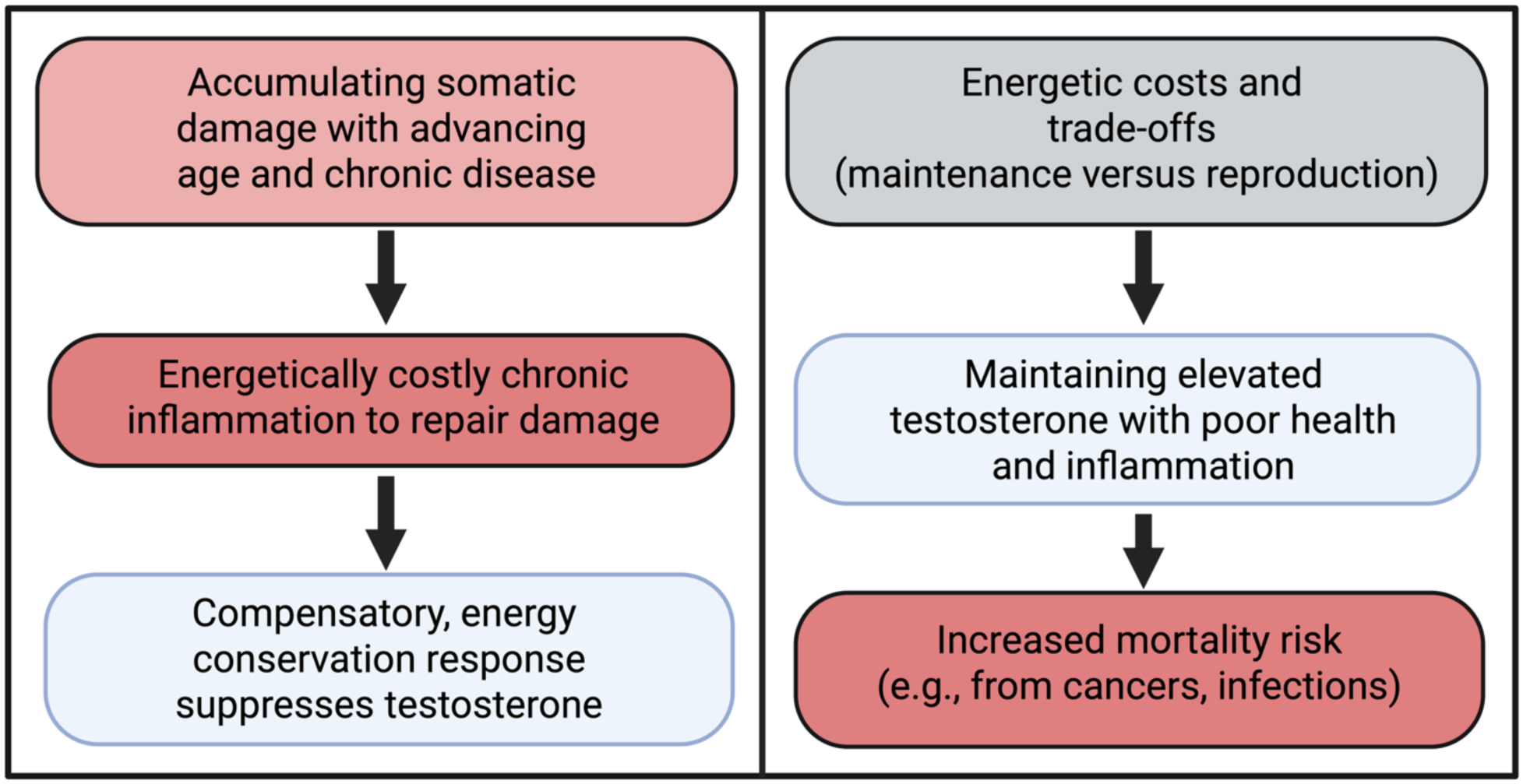

## Introduction

Later life testosterone declines are associated with a variety of health conditions [1–4], as well as low mood and quality of life [5]. Efforts to ameliorate age-related testosterone declines include replacement therapy, which presents concerns regarding long-term safety [6, 7]. Due to great societal interest in attenuating or preventing testosterone declines without introducing new health risks, further research is needed to understand this age-related process.

A recently proposed theory, the Brain-Body Energy Conservation model of aging, suggests that age-related testosterone declines can at least partly be understand as an energy conservation response to developing chronic inflammation [8]. The progressive accumulation of damaged and dysfunctional cells in later life begins to chronically activate the inflammatory response for cellular and tissue repair [9]. This includes the accumulation of senescent cells, which contribute to inflammaging through the senescence-associated secretory phenotype [8, 10, 11]. The inflammatory response requires energy to initiate and sustain, while senescent cells show greater mitochondrial density, indicating greater energy expenditure [8]. However, total body energy expenditure does not increase in later life [12]. This suggests increasing metabolic stress, which is expected to be offset or trade-off with other bodily functions not immediately necessary for survival.

The Brain-Body Energy Conservation model draws from evolutionary life history theory, which seeks to understand how organisms allocate their finite resources, including energy, between competing functions to maximize survival and reproduction [13]. Central to life history theory is the concept of trade-offs, as energy invested in one function comes at the expense of another. These trade-offs are often studied between three broad categories of growth, reproduction, and maintenance, with immune function considered part of maintenance [13–16]. Multiple lines of evidence suggest inflammation is energetically costly and can induce trade-offs with other functions. This includes sickness behaviors, such as fatigue and reduced physical activity [17]. Another commonly studied trade-off is between infection-related immune activation and child growth in high pathogen environments [14, 18, 19].

There is also evidence that inflammation and metabolic stress suppress testosterone. Acute infection, which activates an inflammatory response, decreases testosterone in men [15, 20]. Surgery, which induces acute somatic damage and inflammation, has also been found to induce a temporary reduction in androgens [21, 22]. Relatedly, the metabolic stressors of fasting [23, 24] or extreme physical activity also decrease sex hormones [25]. In contrast, moderate exercise and intermittent fasting, which activate cellular repair mechanisms like autophagy that decrease cellular damage and inflammation, are generally associated with increased sex hormones [25].

The central nervous system provides the necessary flow of information and metabolic regulation to orchestrate an energetic trade-off between chronic inflammation and sex hormone signaling. Pro-inflammatory cytokines and other proteins that signal metabolic stress in the body are detected by the hypothalamus, which can decrease hormone signaling by controlling the pituitary gland [8, 26]. A major signaling protein is Growth Differentiation Factor-15 (GDF-15), which has been labeled a metabokine due to its role in signaling metabolic stress [27, 28]. GDF-15 is produced in response to chronic inflammation, and previous experimental studies have shown that it augments an energy conserving immune strategy of tolerance to pathogens [29, 30]. Circulating concentrations of GDF-15 are monitored by the central nervous system through neurons expressing GDNF family receptor α–like [31], thereby providing a mechanistic link to metabolic regulation. GDF-15 is also a central protein in the senescence-associated secretory phenotype, alongside canonical pro-inflammatory cytokines such as interleukins (IL) 1 and 6, as well as tumor necrosis factors (TNF) and their receptors (TNFR) [8, 10, 11].

Here we test the Brain-Body Energy Conservation model linking inflammaging to later life testosterone declines, relying on a large sample of male participants in the UK BioBank (n = 18,347, ages 40-70, mean age = 57). We test whether inflammaging markers, including IL-1β, IL-6, TNF-α, TNFR1, and GDF-15 mediate inverse associations between age and free and bioavailable serum testosterone. We also test whether these markers mediate inverse associations between chronic diseases, which reflect advanced somatic damage, and testosterone. Finally, since the Brain-Body Energy Conservation model posits that reduced testosterone is an energy conserving adaptation to promote immediate survival against somatic damage, we test whether individuals with elevated testosterone for their given health and inflammation have a higher mortality risk over follow up. Testosterone and inflammaging markers were measured concurrently between 2006-2010, with mortality tracking through 2024.

## Methods

The UKB is a population-based cohort that began with the recruitment of approximately 500,000 individuals aged between 40 and 70 years between 2006 and 2010. It has approval from the North West Multi-center Research Ethics Committee (MREC). Written informed consent was obtained from all participants. Participants self-reported whether they had hypertension, cardiovascular disease, diabetes, or cancer. They also self-reported the type of cancer, here we excluded non-melanoma skin cancer from the cancer designation. The proteomics UKB study initially measured approximately 54,000 samples from the baseline years 2006-2010. Proteomic profiling of blood plasma samples was performed using the antibody-based Olink Explore 3072 PEA [32]. This assay includes binding antibodies to measure 2,923 proteins. Measures reflect relative protein abundance in the sample, which were rank normalized by UKB researchers, resulting in approximately normal distributions. This panel of protein measures included canonical markers of inflammaging, including IL-1β, IL-6, TNF-α, TNFR1, and GDF-15. Serum testosterone was measured using a one-step competitive chemiluminescent immunoassay. ICD-10 codes from electronic medical records were used to determine the primary cause of death.

Free and bioavailable testosterone were calculated with the Vermeulen equation, using serum SHBG and albumin [33]. Free and bioavailable testosterone, as well as cytokines, were standardized (mean = 0, SD = 1). We used linear regression models to test associations between cytokines and testosterone. The first set of models included age, reported diseases, and BMI as predictors, while the second and third set of models progressively added cytokines. We considered mediation if the addition of cytokines to the models attenuated associations between age/diseases and testosterone. Mediation analysis was performed using a quasi-Bayesian approach with 1,000 simulations, implemented in the R package “mediation” [34, 35].

Individuals lost to follow up between baseline and 2024 were excluded (0.3% of the sample). Since low testosterone has been found to predict increased mortality risk, which could be due to confounding with poor health [4], we tested non-linear associations between testosterone and mortality risk before and after adjusting for health using high and low tertiles (<33^rd^ and >66^th^ percentiles). We used logistic regression models to predict the dichotomous outcome of death between baseline and 2024 (yes = 1, no = 0). The first set of models only adjusted for age and time of blood sampling at baseline between 2006-2010. The second set adjusted for having hypertension or CVD, diabetes, or caner, as well as BMI. The final set of models further adjusted for cytokines as additional proxies for health status. The code used in the analysis can be found at: https://github.com/jakearonoff/ukb_inflammaging_testosterone.

## Results

### Markers of inflammaging mediate age and disease-related testosterone declines

Age, diseases (hypertension/cardiovascular disease, Type 2 diabetes, cancer), and BMI were positively associated with IL-6, TNF-α, TNFR1, and GDF-15, while associations for IL-1β were mixed (**Supplementary Table S1**). Age, diabetes, cancer, and BMI were independently inversely associated with both free and bioavailable testosterone (**Table 1**). Having high blood pressure or CVD was inversely associated with free but not bioavailable testosterone. IL-6 was negatively associated with both free and bioavailable testosterone, while its addition to the model attenuated associations for age, diseases, and BMI. Similarly, GDF-15 was negatively associated with both free and bioavailable testosterone, while its addition to the model attenuated associations for age, diseases, and IL-6. Results for the other cytokines were mixed. IL-1β was positively associated with both free and bioavailable testosterone, while TNF-α was not associated. TNFR1 was negatively associated with bioavailable testosterone, and this association was attenuated after adding GDF-15 to the model. However, TNFR1 became positively associated with free testosterone after inclusion of GDF-15.

**Table 1.** Models predicting free and bioavailable testosterone (n = 18,347)

|  | Free Testosterone (z) |  |  | Bioavailable Testosterone (z) |  |  |
| --- | --- | --- | --- | --- | --- | --- |
| Age<br>(years) | -0.036**<br>(0.001) | -0.034**<br>(0.001) | -0.030**<br>(0.001) | -0.041**<br>(0.001) | -0.038**<br>(0.001) | -0.034**<br>(0.001) |
| High BP or<br>CVD (0,1) | -0.046**<br>(0.016) | -0.040*<br>(0.016) | -0.025<br>(0.016) | -0.021<br>(0.015) | -0.006<br>(0.016) | 0.008<br>(0.016) |
| T2 Diabetes<br>(0,1) | -0.110**<br>(0.028) | -0.103**<br>(0.028) | -0.008<br>(0.029) | -0.102**<br>(0.027) | -0.081**<br>(0.027) | 0.009<br>(0.028) |
| Cancer<br>(0,1) | -0.066*<br>(0.031) | -0.055<sup>t</sup><br>(0.031) | -0.045<br>(0.031) | -0.069*<br>(0.031) | -0.046<br>(0.031) | -0.036<br>(0.031) |
| BMI<br>(kg/m<sup>2</sup>) | -0.020**<br>(0.002) | -0.017**<br>(0.002) | -0.018**<br>(0.002) | -0.024**<br>(0.002) | -0.018**<br>(0.002) | -0.019**<br>(0.002) |
| IL-1β<br>(z) | | 0.019<sup>t</sup><br>(0.011) | 0.019<sup>t</sup><br>(0.011) | | 0.024*<br>(0.011) | 0.024*<br>(0.011) |
| IL-6<br>(z) |  | -0.054**<br>(0.009) | -0.038**<br>(0.009) |  | -0.073**<br>(0.009) | -0.057**<br>(0.009) |
| TNF-α<br>(z) | | 0.008<br>(0.020) | 0.027<br>(0.020) | | 0.010<br>(0.020) | 0.028<br>(0.020) |
| TNFR1<br>(z) |  | -0.017<br>(0.023) | 0.095**<br>(0.026) |  | -0.113**<br>(0.023) | -0.006<br>(0.025) |
| GDF-15<br>(z) |  |  | -0.180**<br>(0.017) |  |  | -0.172**<br>(0.016) |

Mean age = 57 years, range 40-70, 36% with high BP or CVD, 8% with Type 2 diabetes, 5% with cancer, mean BMI = 27.8 kg/m<sup>2</sup>, mean free testosterone = 0.21 nmol/L (SD = 0.06), mean bioavailable testosterone = 5.22 (SD = 1.47).
<sup>t</sup>p < 0.10; \* p < 0.05; \*\* p < 0.01.

Significant mediation paths are shown in **Figure 1** and **Supplementary Tables S2-3**. IL-6 and GDF-15 together mediated 17% of the negative association between age and free testosterone, while adjusting for each cytokine showed independent mediation paths of 2% for IL-6 and 12% for GDF-15. Similarly, IL-6 and GDF-15 together mediated 46% of the negative association between having high blood pressure or CVD and free testosterone, while adjusting for each cytokine showed independent mediation paths of 8% for IL-6 and 37% for GDF-15. Finally, IL-6 and GDF-15 together mediated 32% of the negative association between having cancer and free testosterone, while adjusting for each cytokine showed independent mediation paths of 9% for IL-6 and 18% for GDF-15. GDF-15 also mediated 91% of the inverse association between having Type 2 diabetes and free testosterone, while IL-6 mediated 11% of the inverse association between BMI and free testosterone. Finally, GDF-15 mediated 30% of the inverse association between IL-6 and free testosterone.

**Figure 1.**
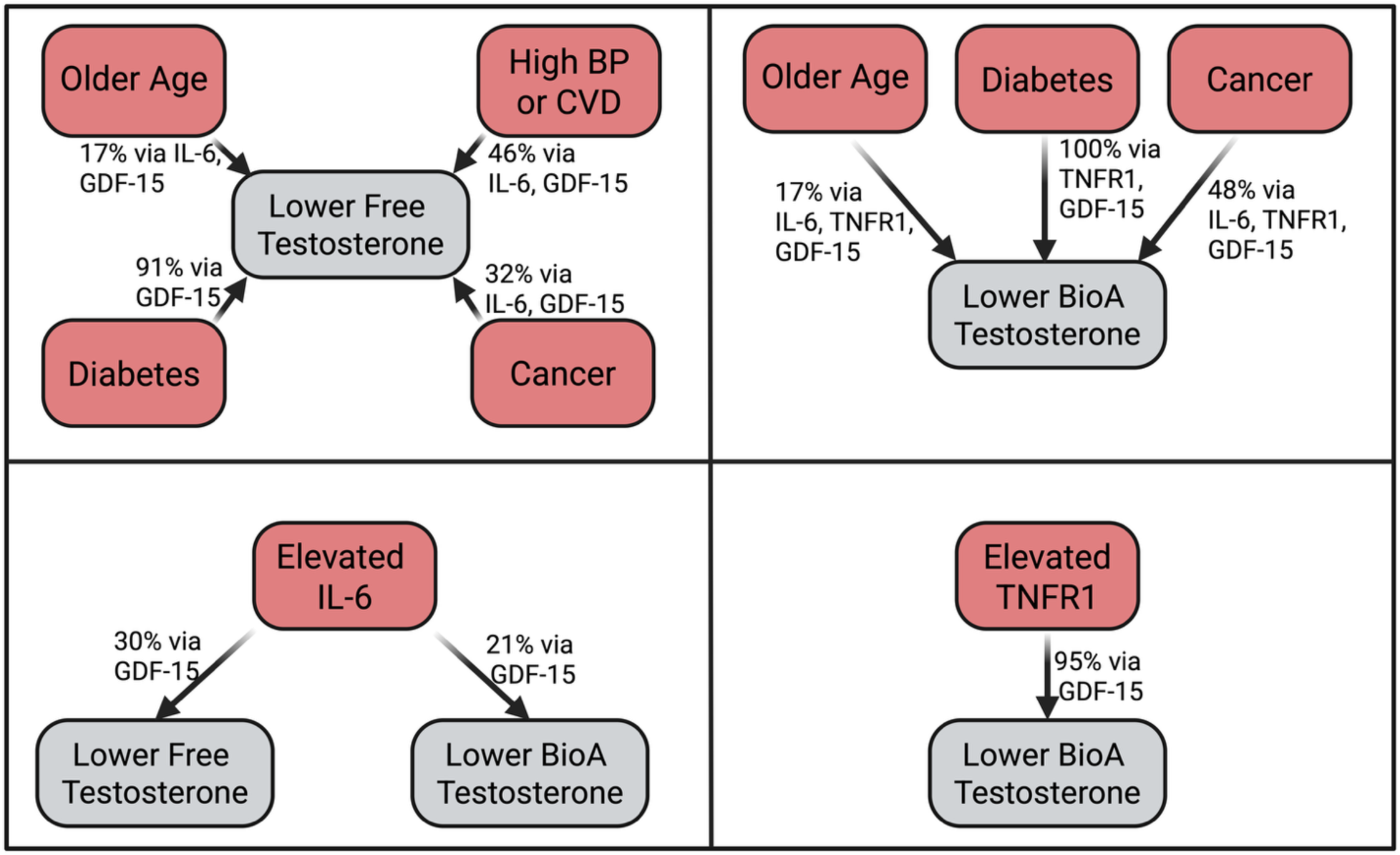
Summary of cytokine mediation results. High BP or CVD refers to hypertension or cardiovascular disease, while diabetes refers to type 2 diabetes and cancer refers to any type excluding non-melanoma skin cancer.

IL-6, TNFR1, and GDF-15 together mediated 17% of the inverse association between age and bioavailable testosterone, while adjusting for each cytokine showed independent mediation paths of 3% for IL-6, 2% for TNFR1, and 11% for GDF-15. TNFR1 and GDF-15 together mediated 100% of the inverse association between diabetes and bioavailable testosterone, while adjusting for each cytokine showed independent mediation paths of 13% for TNFR1 and 100% for GDF-15. IL-6, TNFR1, and GDF-15 together mediated 17% of the inverse association between cancer and bioavailable testosterone, while adjusting for each cytokine showed independent mediation paths of 13% for IL-6, 13% for TNFR1, and 19% for GDF-15. IL-6 and TNFR1 together mediated 25% of the inverse association between BMI and bioavailable testosterone, while adjusting for each cytokine showed independent mediation paths of 13% for IL-6 and 6% for TNFR1. Finally, GDF-15 mediated 21% of the inverse associations between IL-6 and bioavailable testosterone, as well as 5% of the inverse association for TNFR1.

### Maintaining higher testosterone despite poor health increases mortality risk

Before adjusting for diseases, BMI, and inflammation, low free and bioavailable testosterone (<33^rd^ percentile) predicted increased odds of all-cause mortality (**Figure 2**). However, progressively adjusting for these health proxies attenuated the associations. In contrast, high free and bioavailable testosterone (>66^th^ percentile) became predictive of increased odds for all-cause mortality following model inclusion of diseases, BMI, and inflammation. In these fully adjusted models, high free testosterone predicted 12% higher odds of mortality, while high bioavailable testosterone predicted 17% higher odds. For reference, one year of age predicted 9% higher odds of mortality (**Supplementary Table S4**).

**Figure 2.**
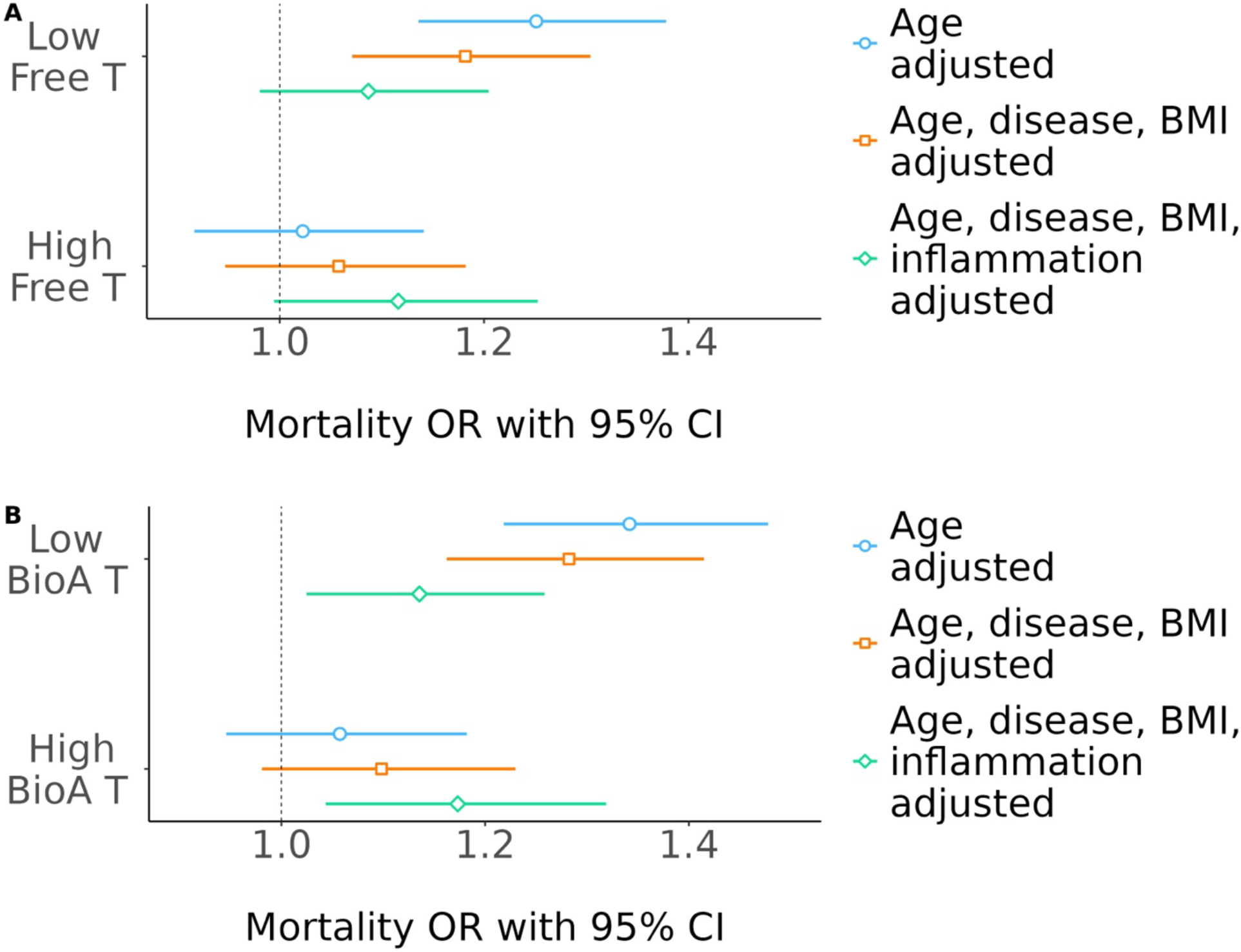
High and low testosterone (**A** = Free T, **B** = Bioavailable T) predicting mortality risk, progressively adjusting for age, diseases, BMI, and cytokines. Models also adjust for time of baseline blood sample (2004-2010).

Considering specific causes of mortality suggested that the all-cause mortality effect is likely attributable to a combination of cancer and infection sources (**Table 2 and Figure 3**). While confidence intervals widened with the smaller number of deaths in cause-specific analyses, the effect size for high free testosterone predicting cancer mortality risk was identical to all-causes, at 12% higher odds. For infection-specific deaths, high free testosterone predicted 26% higher odds, while high bioavailable testosterone predicted 27% higher odds. For cancer and infection deaths combined, high free testosterone predicted 14% higher odds, while high bioavailable testosterone predicted 15% higher odds. For reference, one year of age predicted 7% higher odds of cancer or infection mortality (**Supplementary Table S4**). Effect sizes for high testosterone predicting cardiovascular-specific death were weaker and inconsistent.

**Figure 3.**
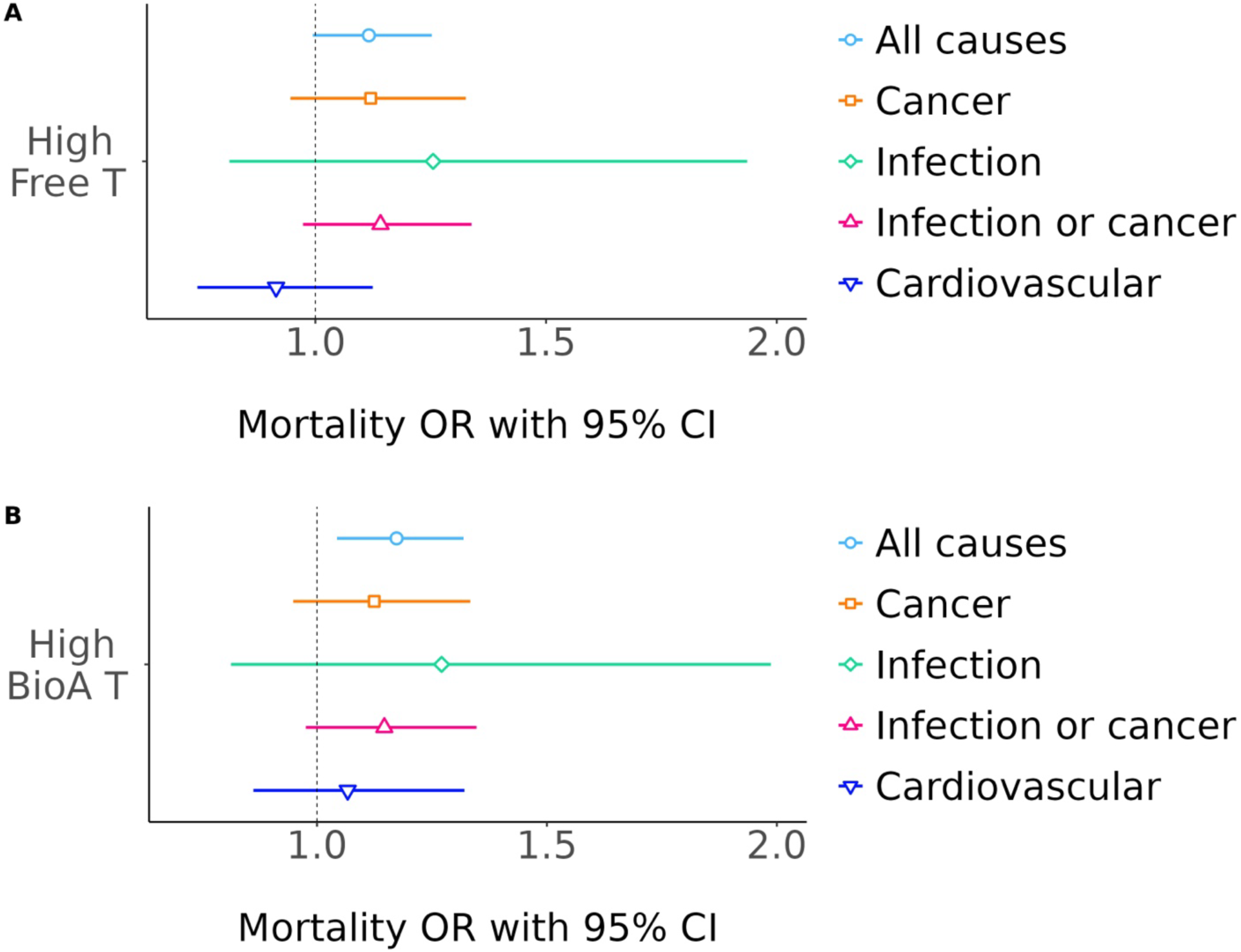
High testosterone (**A** = Free T, **B** = Bioavailable T) predicting mortality risk through 2024, adjusting for time of baseline blood sample (2004-2010), age, diseases, BMI, and cytokines.

**Table 2.** Testosterone at baseline (2006-2010, n = 18,347) predicting mortality through 2024. Estimates are odds ratios with 95% CI from logistic regression models.

|  | All causes<br>(3,073 deaths) |  | Cancer<br>(1,090 deaths) |  | Infection<br>(153 deaths) |  | Cancer or<br>Infection<br>(1,243 deaths) |  | Cardiovascular<br>(763 deaths) |  |
| --- | --- | --- | --- | --- | --- | --- | --- | --- | --- | --- |
|  | Free | BioA | Free | BioA | Free | BioA | Free | BioA | Free | BioA |
| Low T | 1.09<br>(0.98,<br>1.20) |  | 1.10<br>(0.95,<br>1.28) |  | 0.95<br>(0.65,<br>1.41) |  | 1.08<br>(0.94,<br>1.25) |  | 0.90<br>(0.75,<br>1.07) |  |
| High T | 1.12<sup>†</sup><br>(0.99,<br>1.25) |  | 1.12<br>(0.95,<br>1.33) |  | 1.26<br>(0.81,<br>1.94) |  | 1.14<br>(0.97,<br>1.34) |  | 0.91<br>(0.74,<br>1.12) |  |
| Low T |  | 1.14*<br>(1.02,<br>1.26) |  | 1.05<br>(0.91,<br>1.23) |  | 1.01<br>(0.69,<br>1.49) |  | 1.05<br>(0.91,<br>1.21) |  | 1.10<br>(0.92,<br>1.31) |
| High T |  | 1.17**<br>(1.04,<br>1.32) |  | 1.12<br>(0.95,<br>1.33) |  | 1.27<br>(0.81,<br>1.98) |  | 1.15<sup>†</sup><br>(0.98,<br>1.35) |  | 1.07<br>(0.86,<br>1.32) |

Reference groups are the middle tertiles of testosterone.
Adjusted for time of blood sample at baseline, age, high BP or CVD, diabetes, cancer, BMI, and cytokines.
Full results with all covariates are shown in Supplementary Tables S4-5.
<sup>†</sup>p < 0.10; \* p < 0.05; \*\* p < 0.01

We also considered deaths specific to prostate cancer (n = 126 deaths), which is augmented by testosterone exposure, versus cancer deaths from other sources (n = 964 deaths) (**Supplementary Table S5**). Results suggested that prostate cancer was a primary, but not exclusive, contributor to the effect for cancer mortality broadly. High free testosterone predicted 21% higher odds of prostate cancer death and 10% higher odds of death from cancers other than prostate, while high bioavailable testosterone predicted 11% higher odds of prostate cancer death and 12% higher odds of death from cancers other than prostate.

## Discussion

Here we find evidence consistent with the Brain-Body Energy Conservation model of aging, in which testosterone declines in later life are partly due to an energetic trade-off with inflammation. GDF-15, a central marker of the senescence-associated secretory phenotype as well as metabolic stress, was inversely associated with testosterone and a key mediator in the inverse associations with age and chronic disease. Our results are consistent with previous studies identifying GDF-15 as one of the strongest predictors of aging and health: one standard deviation increase had an equivalent effect on free testosterone as being five years older in age. Further, these results are consistent with our previous study reporting evidence for an energy conservation model of immunosenescence [36]. Finally, consistent with a trade-off between reproduction and maintenance, having high testosterone adjusted for health and inflammation came with the expected cost of increased mortality risk. While statistical power was limited to decipher specific mortality causes, there was suggestive evidence this effect was attributable to a combination of cancer, particularly prostate, and infection mortality.

This study has important health and therapeutic implications. It highlights targeting cellular damage that elicits inflammation to alleviate later life testosterone declines [37]. This can be achieved through lifestyle factors like moderate caloric restriction and increasing physical activity. It might also be achieved with therapeutics that target later life inflammation. For example, rapamycin inhibits mTOR, leading to activation of AMPK and downstream cellular maintenance mechanisms like autophagy that reduce age-related damage and inflammation [38, 39]. Other therapeutics like metformin and GLP-1 receptor agonists achieve similar outcomes related to increased cellular repair and reduced inflammation, potentially alleviating testosterone declines [37, 40, 41]. While GDF-15 was inversely associated with testosterone, this finding does not suggest that therapeutically suppressing it is an appropriate strategy. GDF-15 is likely making the best of a bad situation, concentrating energy expenditure on the inflammatory response to repair cellular and tissue damage [42]. Inhibiting GDF-15 might elevate testosterone declines at the cost of this necessary maintenance, leading to short-term benefits at the expense of long-term health [8]. Finally, our results of elevated testosterone and mortality risk emphasize potential risks of exogenous testosterone administration without addressing the underlying inflammation.

Similar to what has been reported elsewhere, low testosterone also predicted increased mortality risk, and the effect was attenuated after adjusting for cytokines [4, 43]. This is likely due to confounding with poor health, which can cause both low testosterone and increased mortality risk. Our results showing progressive attenuation of the low testosterone and mortality risk effect after adjusting for health indicators supports this explanation. While the association between low testosterone and mortality risk was not completely attenuated in these fully adjusted models, this is likely attributable to residual confounding. Specifically, an individual’s somatic damage could be worse than their age, reported disease, and single time point inflammation measures indicate. This would result in having lower testosterone after adjusting for these covariates as well as an increased mortality risk. Similarly, some individuals likely have better health than indicated by self-reports and a single measure of inflammation, creating further residual confounding. Our results are therefore likely underestimating the true effect of higher testosterone on mortality risk.

Associations varied in magnitude across testosterone and inflammatory measures, for example they tended to be stronger for bioavailable versus free testosterone. This might be due to the greater concentration and variability of bioavailable testosterone, resulting in improved statistical power. For cytokines, the metabokine GDF-15 was the strongest predictor of testosterone. This could be suggesting the central importance of energetic stress in response to cellular damage as a contributor to testosterone declines, in contrast to inflammation in general.

Further evidence that chronic inflammation in later life is an important contributor to testosterone declines comes from comparisons with non-industrialized human populations, such as Tsimane forager-farmers in the Bolivian Amazon. The Tsimane live a highly active lifestyle with minimal excess caloric consumption [44]. They have the lowest reported prevalence of cardiometabolic disease [45, 46]. Inflammaging is markedly attenuated in the Tsimane [47–49]. Consistent with the Brain-Body Energy Conservation model, age-related testosterone declines are attenuated in the Tsimane [50], and we do not find that low testosterone is associated with CVD [51]. Replicating our findings in a subsistence population like the Tsimane would strengthen evidence for the Brain-Body Energy Conservation model of aging.

Our study highlights the importance of considering energetic trade-offs from a life history perspective, as outlined in the Brain-Body Energy Conservation model of aging, to understand functional declines in later life. Here we find evidence suggesting that metabolic stress from accumulating somatic damage and inflammation leads to a trade-off with testosterone. This trade-off between reproduction and maintenance is further highlighted by the finding of high testosterone predicting increased mortality risk. Future studies can help clarify whether this trade-off model explains other functional declines as well as tissue atrophy in later life.

## Acknowledgements

This research has been conducted using the UK Biobank Resource under the approved application number 772113. We are grateful to the UKB study participants, as well as the researchers who have created and maintained the UKB data. Figure 1 and the graphical abstract were created in Biorender. Funded in part by the National Institutes of Health (NIH) National Institute on Aging (NIA) (R01AG054442).

## Data availability

UK Biobank data are available to researchers with approved project applications and payment of applicable access costs.

## Author contributions

Jacob E. Aronoff: conceptualization (lead), statistical analysis, writing (original draft). Benjamin C. Trumble: conceptualization (supporting), writing (review and editing).

---

**Supplementary Table S1.** Models predicting cytokines (n = 18,347)

| | IL-1β (z) | IL-6 (z) | TNF-α (z) | TNFR1 (z) | GDF-15 (z) |
| --- | --- | --- | --- | --- | --- |
| Age<br>(years) | 0.001<sup>t</sup><br>(0.001) | 0.020**<br>(0.001) | 0.005**<br>(0.0004) | 0.010**<br>(0.0003) | 0.032**<br>(0.0005) |
| High BP or<br>CVD<br>(0,1) | 0.032**<br>(0.011) | 0.120**<br>(0.014) | 0.054**<br>(0.006) | 0.068**<br>(0.006) | 0.141**<br>(0.008) |
| Diabetes<br>(0,1) | -0.033<sup>t</sup><br>(0.018) | 0.093**<br>(0.024) | 0.047**<br>(0.011) | 0.123**<br>(0.010) | 0.615**<br>(0.014) |
| Cancer<br>(0,1) | -0.037<sup>t</sup><br>(0.021) | 0.165**<br>(0.028) | 0.049**<br>(0.013) | 0.091**<br>(0.011) | 0.133**<br>(0.016) |
| BMI<br>(kg/m<sup>2</sup>) | 0.008**<br>(0.001) | 0.051**<br>(0.002) | 0.012**<br>(0.001) | 0.019**<br>(0.001) | 0.015**<br>(0.001) |

<sup>t</sup> p < 0.10; \* p < 0.05; \*\* p < 0.01

**Supplementary Table S2.** Mediation results for age, diseases, cytokines, and testosterone in men in the UKB (n = 18,347)

| Predictor | Mediator | Outcome | Proportion mediated |
| --- | --- | --- | --- |
| Age | IL-6 | Free T | 0.02 |
| High BP or CVD | IL-6 | Free T | 0.08 |
| Cancer | IL-6 | Free T | 0.09 |
| BMI | IL-6 | Free T | 0.11 |
| Age | GDF-15 | Free T | 0.12 |
| High BP or CVD | GDF-15 | Free T | 0.37 |
| Diabetes | GDF-15 | Free T | 0.91 |
| Cancer | GDF-15 | Free T | 0.18 |
| IL-6 | GDF-15 | Free T | 0.30 |

**Supplementary Table S3.**
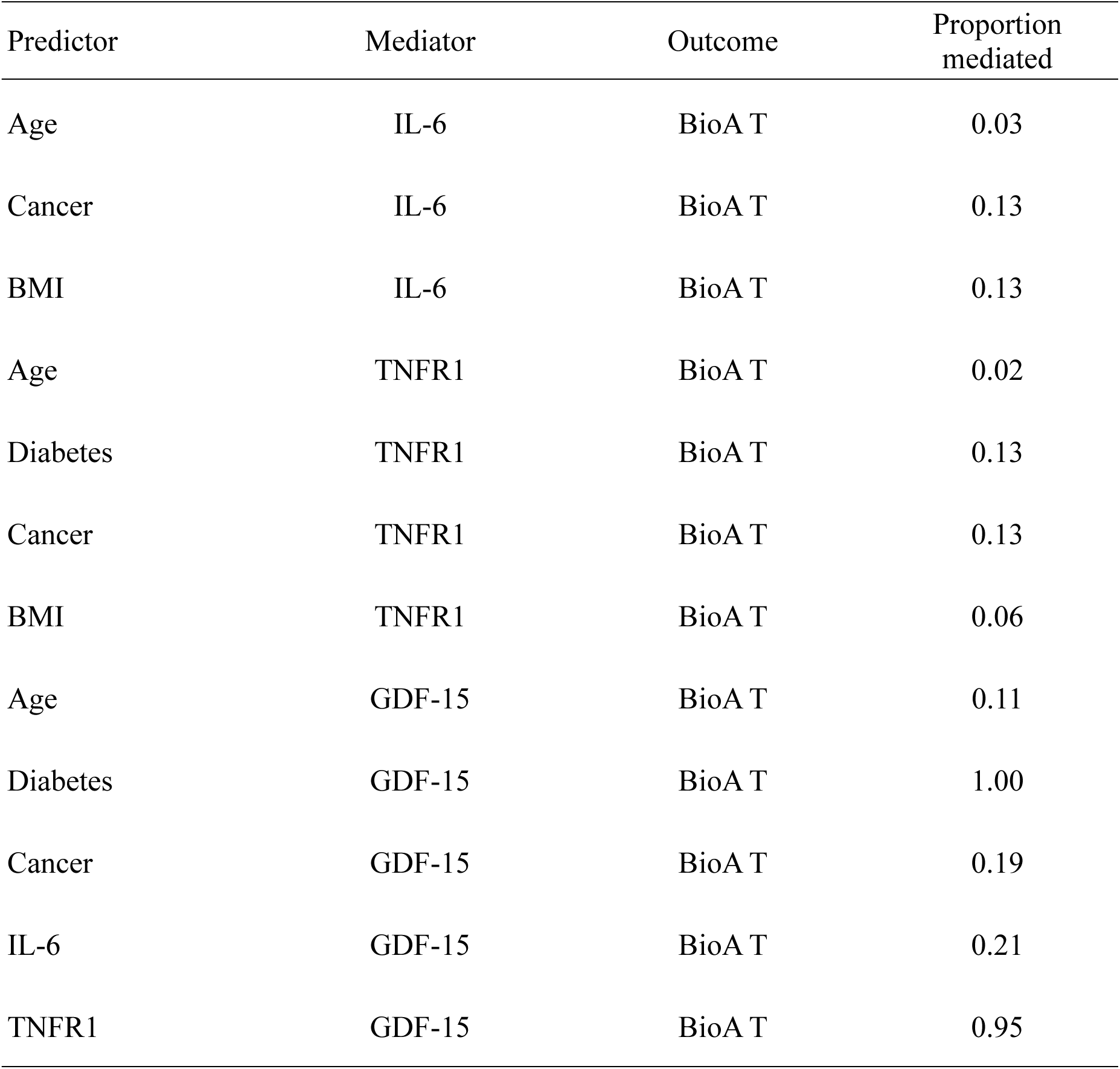
Mediation results for age, diseases, cytokines, and testosterone in men in the UKB (n = 18,347)

| Predictor | Mediator | Outcome | Proportion mediated |
| --- | --- | --- | --- |
| Age | IL-6 | BioA T | 0.03 |
| Cancer | IL-6 | BioA T | 0.13 |
| BMI | IL-6 | BioA T | 0.13 |
| Age | TNFR1 | BioA T | 0.02 |
| Diabetes | TNFR1 | BioA T | 0.13 |
| Cancer | TNFR1 | BioA T | 0.13 |
| BMI | TNFR1 | BioA T | 0.06 |
| Age | GDF-15 | BioA T | 0.11 |
| Diabetes | GDF-15 | BioA T | 1.00 |
| Cancer | GDF-15 | BioA T | 0.19 |
| IL-6 | GDF-15 | BioA T | 0.21 |
| TNFR1 | GDF-15 | BioA T | 0.95 |

**Supplementary Table S4.** Logistic regression models predicting mortality (OR with 95% CI) (n = 18,347)

|  | All causes<br>(n = 3,073 deaths) |  | Cancer<br>(n = 1,090 deaths) |  | Infection<br>(n = 153 deaths) |  | Cancer or Infection<br>(n = 1,243 deaths) |  |
| --- | --- | --- | --- | --- | --- | --- | --- | --- |
| Time<br>(years) | 0.76**<br>(0.72,<br>0.80) | 0.76**<br>(0.72,<br>0.80) | 0.86**<br>(0.80,<br>0.93) | 0.86**<br>(0.80,<br>0.93) | 0.94<br>(0.77,<br>1.14) | 0.94<br>(0.77,<br>1.14) | 0.87**<br>(0.81,<br>0.94) | 0.87**<br>(0.81,<br>0.93) |
| Age<br>(years) | 1.09**<br>(1.08,<br>1.10) | 1.09**<br>(1.08,<br>1.10) | 1.07**<br>(1.06,<br>1.08) | 1.07**<br>(1.06,<br>1.08) | 1.07**<br>(1.04,<br>1.11) | 1.07**<br>(1.04,<br>1.11) | 1.07**<br>(1.06,<br>1.08) | 1.07**<br>(1.06,<br>1.08) |
| High BP or<br>CVD (0,1) | 1.29**<br>(1.18,<br>1.41) | 1.29**<br>(1.18,<br>1.42) | 1.03<br>(0.90,<br>1.17) | 1.03<br>(0.90,<br>1.18) | 1.46*<br>(1.03,<br>2.10) | 1.46*<br>(1.03,<br>2.10) | 1.07<br>(0.94,<br>1.22) | 1.07<br>(0.94,<br>1.22) |
| Diabetes<br>(0,1) | 1.06<br>(0.91,<br>1.23) | 1.06<br>(0.91,<br>1.23) | 0.80*<br>(0.63,<br>1.00) | 0.80<sup>†</sup><br>(0.63,<br>1.00) | 1.28<br>(0.80,<br>2.01) | 1.27<br>(0.79,<br>2.00) | 0.87<br>(0.71,<br>1.07) | 0.87<br>(0.71,<br>1.07) |
| Cancer<br>(0,1) | 1.67**<br>(1.43,<br>1.96) | 1.67**<br>(1.43,<br>1.96) | 2.50**<br>(2.07,<br>3.01) | 2.50**<br>(2.07,<br>3.01) | 1.08<br>(0.59,<br>1.84) | 1.08<br>(0.59,<br>1.84) | 2.35**<br>(1.95,<br>2.81) | 2.35**<br>(1.95,<br>2.81) |
| BMI<br>(kg/m<sup>2</sup>) | 0.99*<br>(0.98,<br>1.00) | 0.99*<br>(0.98,<br>1.00) | 0.99<br>(0.97,<br>1.01) | 0.99<br>(0.97,<br>1.01) | 1.02<br>(0.99,<br>1.06) | 1.02<br>(0.99,<br>1.06) | 0.99<br>(0.98,<br>1.01) | 0.99<br>(0.98,<br>1.01) |
| IL-1β<br>(z) | 1.09**<br>(1.02,<br>1.17) | 1.09*<br>(1.02,<br>1.17) | 1.12*<br>(1.02,<br>1.24) | 1.12*<br>(1.02,<br>1.24) | 0.84<br>(0.64,<br>1.09) | 0.84<br>(0.64,<br>1.09) | 1.08<sup>†</sup><br>(0.99,<br>1.19) | 1.08<sup>†</sup><br>(0.99,<br>1.19) |
| IL-6<br>(z) | 1.23**<br>(1.17,<br>1.29) | 1.22**<br>(1.16,<br>1.29) | 1.16**<br>(1.08,<br>1.24) | 1.16**<br>(1.08,<br>1.24) | 1.27**<br>(1.08,<br>1.46) | 1.27**<br>(1.08,<br>1.46) | 1.18**<br>(1.11,<br>1.26) | 1.18**<br>(1.11,<br>1.26) |
| TNF-α<br>(z) | 1.01<br>(0.89,<br>1.14) | 1.01<br>(0.90,<br>1.15) | 0.94<br>(0.78,<br>1.12) | 0.94<br>(0.78,<br>1.12) | 0.92<br>(0.57,<br>1.40) | 0.92<br>(0.57,<br>1.40) | 0.93<br>(0.78,<br>1.10) | 0.93<br>(0.78,<br>1.10) |
| TNFR1<br>(z) | 1.06<br>(0.92,<br>1.23) | 1.06<br>(0.91,<br>1.23) | 0.91<br>(0.74,<br>1.12) | 0.91<br>(0.74,<br>1.12) | 1.22<br>(0.76,<br>1.94) | 1.22<br>(0.76,<br>1.94) | 0.96<br>(0.79,<br>1.16) | 0.95<br>(0.78,<br>1.16) |
| GDF-15<br>(z) | 2.96**<br>(2.69,<br>3.27) | 2.96**<br>(2.69,<br>3.26) | 1.89**<br>(1.66,<br>2.14) | 1.89**<br>(1.67,<br>2.15) | 1.85**<br>(1.38,<br>2.43) | 1.84**<br>(1.38,<br>2.42) | 1.94**<br>(1.72,<br>2.18) | 1.94**<br>(1.72,<br>2.19) |
| Low Free T | 1.09<br>(0.98,<br>1.20) |  | 1.10<br>(0.95,<br>1.28) |  | 0.95<br>(0.65,<br>1.41) |  | 1.08<br>(0.94,<br>1.25) |  |
| High Free T | 1.12<sup>†</sup><br>(0.99,<br>1.25) |  | 1.12<br>(0.95,<br>1.33) |  | 1.26<br>(0.81,<br>1.94) |  | 1.14<br>(0.97,<br>1.34) |  |
| Low BioA T |  | 1.14*<br>(1.02,<br>1.26) |  | 1.05<br>(0.91,<br>1.23) |  | 1.01<br>(0.69,<br>1.49) |  | 1.05<br>(0.91,<br>1.21) |
| High BioA T |  | 1.17**<br>(1.04,<br>1.32) |  | 1.12<br>(0.95,<br>1.33) |  | 1.27<br>(0.81,<br>1.98) |  | 1.15<sup>†</sup><br>(0.98,<br>1.35) |

<sup>†</sup>p < 0.10; \* p < 0.05; \*\* p < 0.01

**Supplementary Table S5.**
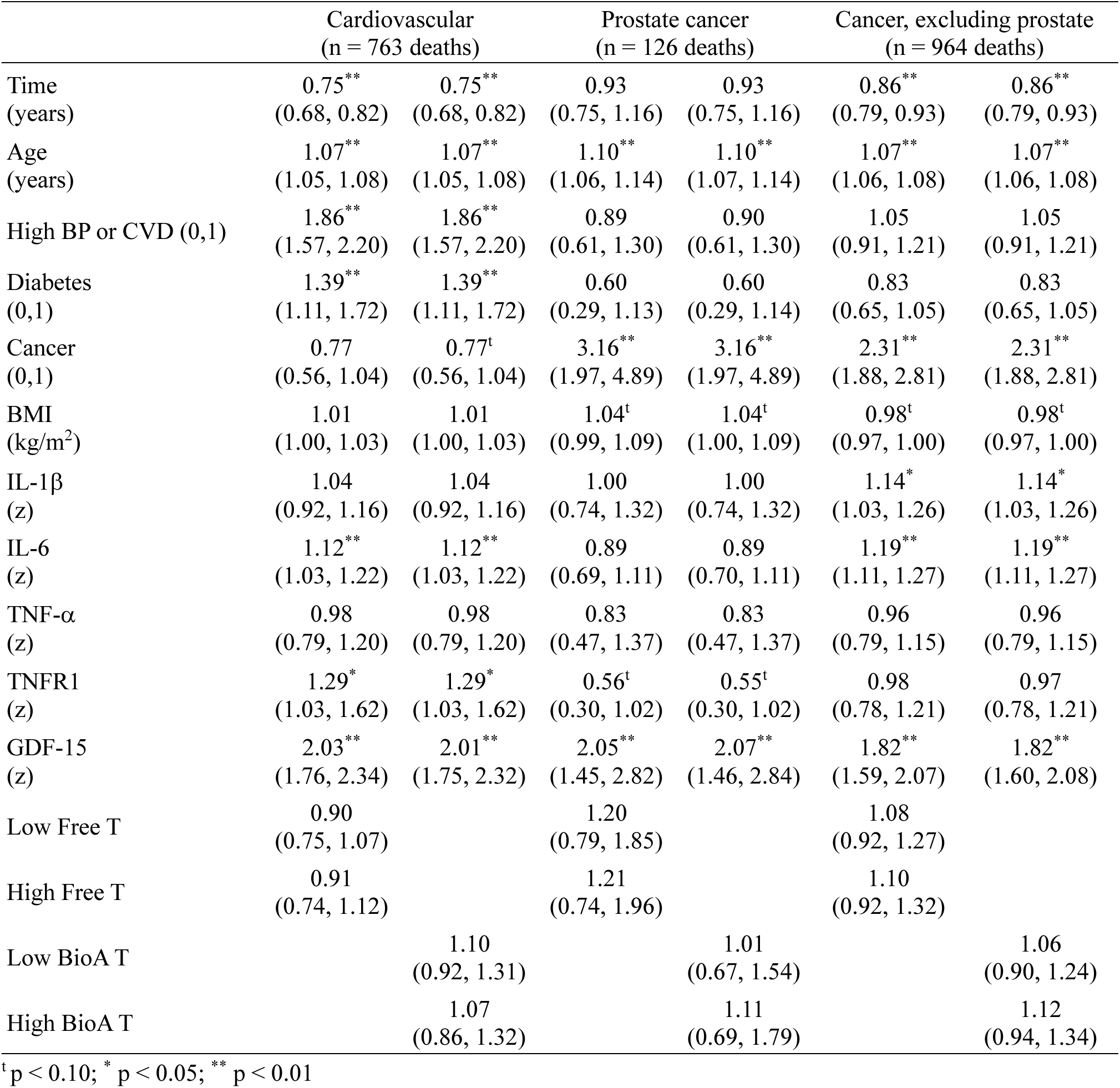
Logistic regression models predicting mortality (OR with 95% CI) (n = 18,347)

|  | Cardiovascular<br>(n = 763 deaths) |  | Prostate cancer<br>(n = 126 deaths) |  | Cancer, excluding prostate<br>(n = 964 deaths) |  |
| --- | --- | --- | --- | --- | --- | --- |
| Time<br>(years) | 0.75**<br>(0.68, 0.82) | 0.75**<br>(0.68, 0.82) | 0.93<br>(0.75, 1.16) | 0.93<br>(0.75, 1.16) | 0.86**<br>(0.79, 0.93) | 0.86**<br>(0.79, 0.93) |
| Age<br>(years) | 1.07**<br>(1.05, 1.08) | 1.07**<br>(1.05, 1.08) | 1.10**<br>(1.06, 1.14) | 1.10**<br>(1.07, 1.14) | 1.07**<br>(1.06, 1.08) | 1.07**<br>(1.06, 1.08) |
| High BP or CVD (0,1) | 1.86**<br>(1.57, 2.20) | 1.86**<br>(1.57, 2.20) | 0.89<br>(0.61, 1.30) | 0.90<br>(0.61, 1.30) | 1.05<br>(0.91, 1.21) | 1.05<br>(0.91, 1.21) |
| Diabetes<br>(0,1) | 1.39**<br>(1.11, 1.72) | 1.39**<br>(1.11, 1.72) | 0.60<br>(0.29, 1.13) | 0.60<br>(0.29, 1.14) | 0.83<br>(0.65, 1.05) | 0.83<br>(0.65, 1.05) |
| Cancer<br>(0,1) | 0.77<br>(0.56, 1.04) | 0.77<sup>t</sup><br>(0.56, 1.04) | 3.16**<br>(1.97, 4.89) | 3.16**<br>(1.97, 4.89) | 2.31**<br>(1.88, 2.81) | 2.31**<br>(1.88, 2.81) |
| BMI<br>(kg/m<sup>2</sup>) | 1.01<br>(1.00, 1.03) | 1.01<br>(1.00, 1.03) | 1.04<sup>t</sup><br>(0.99, 1.09) | 1.04<sup>t</sup><br>(1.00, 1.09) | 0.98<sup>t</sup><br>(0.97, 1.00) | 0.98<sup>t</sup><br>(0.97, 1.00) |
| IL-1β<br>(z) | 1.04<br>(0.92, 1.16) | 1.04<br>(0.92, 1.16) | 1.00<br>(0.74, 1.32) | 1.00<br>(0.74, 1.32) | 1.14*<br>(1.03, 1.26) | 1.14*<br>(1.03, 1.26) |
| IL-6<br>(z) | 1.12**<br>(1.03, 1.22) | 1.12**<br>(1.03, 1.22) | 0.89<br>(0.69, 1.11) | 0.89<br>(0.70, 1.11) | 1.19**<br>(1.11, 1.27) | 1.19**<br>(1.11, 1.27) |
| TNF-α<br>(z) | 0.98<br>(0.79, 1.20) | 0.98<br>(0.79, 1.20) | 0.83<br>(0.47, 1.37) | 0.83<br>(0.47, 1.37) | 0.96<br>(0.79, 1.15) | 0.96<br>(0.79, 1.15) |
| TNFR1<br>(z) | 1.29*<br>(1.03, 1.62) | 1.29*<br>(1.03, 1.62) | 0.56<sup>t</sup><br>(0.30, 1.02) | 0.55<sup>t</sup><br>(0.30, 1.02) | 0.98<br>(0.78, 1.21) | 0.97<br>(0.78, 1.21) |
| GDF-15<br>(z) | 2.03**<br>(1.76, 2.34) | 2.01**<br>(1.75, 2.32) | 2.05**<br>(1.45, 2.82) | 2.07**<br>(1.46, 2.84) | 1.82**<br>(1.59, 2.07) | 1.82**<br>(1.60, 2.08) |
| Low Free T | 0.90<br>(0.75, 1.07) |  | 1.20<br>(0.79, 1.85) |  | 1.08<br>(0.92, 1.27) |  |
| High Free T | 0.91<br>(0.74, 1.12) |  | 1.21<br>(0.74, 1.96) |  | 1.10<br>(0.92, 1.32) |  |
| Low BioA T |  | 1.10<br>(0.92, 1.31) |  | 1.01<br>(0.67, 1.54) |  | 1.06<br>(0.90, 1.24) |
| High BioA T |  | 1.07<br>(0.86, 1.32) |  | 1.11<br>(0.69, 1.79) |  | 1.12<br>(0.94, 1.34) |

<sup>t</sup>p < 0.10; \* p < 0.05; \*\* p < 0.01

## References

1. Khaw, K.-T., et al. Endogenous testosterone and mortality due to all causes, cardiovascular disease, and cancer in men: European prospective investigation into cancer in Norfolk (EPIC-Norfolk) Prospective Population Study. Circulation. 2007;116:2694–2701. 10.1161/CIRCULATIONAHA.107.719005.

2. Ohlsson, C., et al. High serum testosterone is associated with reduced risk of cardiovascular events in elderly men: the MrOS (Osteoporotic Fractures in Men) study in Sweden. Journal of the American College of Cardiology. 2011;58:1674–1681. 10.1016/j.jacc.2011.07.019.

3. Gettler, L.T., et al. Adiposity, CVD risk factors and testosterone: Variation by partnering status and residence with children in US men. Evol Med Public Health. 2017;2017:67–80. 10.1093/emph/eox005.

4. Laughlin, G.A., E. Barrett-Connor, and J. Bergstrom. Low serum testosterone and mortality in older men. J Clin Endocrinol Metab. 2008;93:68–75. 10.1210/jc.2007-1792.

5. Biagetti, B. and M. Puig-Domingo. Age-Related Hormones Changes and Its Impact on Health Status and Lifespan. Aging Dis. 2023;14:605–620. 10.14336/AD.2022.1109.

6. Cheng, H., et al. Age-related testosterone decline: mechanisms and intervention strategies. Reprod Biol Endocrinol. 2024;22:144. 10.1186/s12958-024-01316-5.

7. Rodrigues Dos Santos, M. and S. Bhasin. Benefits and Risks of Testosterone Treatment in Men with Age-Related Decline in Testosterone. Annu Rev Med. 2021;72:75–91. 10.1146/annurev-med-050219-034711.

8. Shaulson, E.D., A.A. Cohen, and M. Picard. The brain-body energy conservation model of aging. Nat Aging. 2024;4:1354–1371. 10.1038/s43587-024-00716-x.

9. Franceschi, C. and J. Campisi. Chronic inflammation (inflammaging) and its potential contribution to age-associated diseases. J Gerontol A Biol Sci Med Sci. 2014;69 Suppl 1:S4–9. 10.1093/gerona/glu057.

10. Coppe, J.P., et al. Senescence-associated secretory phenotypes reveal cell-nonautonomous functions of oncogenic RAS and the p53 tumor suppressor. PLoS Biol. 2008;6:2853–68. 10.1371/journal.pbio.0060301.

11. Basisty, N., et al. A proteomic atlas of senescence-associated secretomes for aging biomarker development. PLoS Biol. 2020;18:e3000599. 10.1371/journal.pbio.3000599.

12. Pontzer, H., et al. Daily energy expenditure through the human life course. Science. 2021;373:808–812. 10.1126/science.abe5017.

13. McDade, T.W. Life history theory and the immune system: steps toward a human ecological immunology. Am J Phys Anthropol. 2003;Suppl 37:100–25. 10.1002/ajpa.10398.

14. Urlacher, S.S., et al. Tradeoffs between immune function and childhood growth among Amazonian forager-horticulturalists. Proceedings of the National Academy of Sciences. 2018;115:E3914–E3921. 10.1073/pnas.1717522115.

15. Muehlenbein, M.P., et al. Toward quantifying the usage costs of human immunity: Altered metabolic rates and hormone levels during acute immune activation in men. Am J Hum Biol. 2010;22:546–56. 10.1002/ajhb.21045.

16. Muehlenbein, M.P. and R.G. Bribiescas. Testosterone-mediated immune functions and male life histories. Am J Hum Biol. 2005;17:527–58. 10.1002/ajhb.20419.

17. Shattuck, E.C. and M.P. Muehlenbein. Human sickness behavior: Ultimate and proximate explanations. Am J Phys Anthropol. 2015;157:1–18. 10.1002/ajpa.22698.

18. Garcia, A.R., et al. Evidence for height and immune function trade-offs among preadolescents in a high pathogen population. Evolution, medicine, and public health. 2020;2020:86–99. 10.1093/emph/eoaa017.

19. McDade, T.W., et al. Maintenance versus growth: investigating the costs of immune activation among children in lowland Bolivia. American Journal of Physical Anthropology: The Official Publication of the American Association of Physical Anthropologists. 2008;136:478–484. 10.1002/ajpa.20831.

20. Spratt, D., et al. Reproductive axis suppression in acute illness is related to disease severity. The Journal of Clinical Endocrinology & Metabolism. 1993;76:1548–1554. 10.1210/jcem.76.6.8501163.

21. Burton, D., G. Nicholson, and G. Hall. Endocrine and metabolic response to surgery. Continuing Education in Anaesthesia, Critical Care & Pain. 2004;4:144–147. 10.1093/bjaceaccp/mkh046.

22. Spratt, D.I., et al. Characterization of a prospective human model for study of the reproductive hormone responses to major illness. Am J Physiol Endocrinol Metab. 2008;295:E63–9. 10.1152/ajpendo.00472.2007.

23. Trumble, B.C., et al. Responsiveness of the reproductive axis to a single missed evening meal in young adult males. Am J Hum Biol. 2010;22:775–81. 10.1002/ajhb.21079.

24. Cameron, J.L., et al. Slowing of pulsatile luteinizing hormone secretion in men after forty-eight hours of fasting. J Clin Endocrinol Metab. 1991;73:35–41. 10.1210/jcem-73-1-35.

25. Mennitti, C., et al. How does physical activity modulate hormone responses? Biomolecules. 2024;14:1418. 10.3390/biom14111418.

26. Jones, T.H. and R.L. Kennedy. Cytokines and hypothalamic-pituitary function. Cytokine. 1993;5:531–8. 10.1016/s1043-4666(05)80001-8.

27. Breit, S.N., D.A. Brown, and V.W. Tsai. The GDF15-GFRAL Pathway in Health and Metabolic Disease: Friend or Foe? Annu Rev Physiol. 2021;83:127–151. 10.1146/annurev-physiol-022020-045449.

28. Salminen, A. GDF15/MIC-1: A stress-induced immunosuppressive factor which promotes the aging process. Biogerontology. 2025;26:19. 10.1007/s10522-024-10164-0.

29. Luan, H.H., et al. GDF15 is an inflammation-induced central mediator of tissue tolerance. Cell. 2019;178:1231–1244. e11. 10.1016/j.cell.2019.07.033.

30. Reyes, J. and G.S. Yap. Emerging Roles of Growth Differentiation Factor 15 in Immunoregulation and Pathogenesis. J Immunol. 2023;210:5–11. 10.4049/jimmunol.2200641.

31. Mullican, S.E., et al. GFRAL is the receptor for GDF15 and the ligand promotes weight loss in mice and nonhuman primates. Nature medicine. 2017;23:1150–1157. 10.1038/nm.4392.

32. Sun, B.B., et al. Plasma proteomic associations with genetics and health in the UK Biobank. Nature. 2023;622:329–338. 10.1038/s41586-023-06592-6.

33. Vermeulen, A., L. Verdonck, and J.M. Kaufman. A critical evaluation of simple methods for the estimation of free testosterone in serum. J Clin Endocrinol Metab. 1999;84:3666–72. 10.1210/jcem.84.10.6079.

34. Imai, K., L. Keele, and D. Tingley. A general approach to causal mediation analysis. Psychol Methods. 2010;15:309–34. 10.1037/a0020761.

35. Tingley, D., et al. Mediation: R package for causal mediation analysis. 2014;10.18637/jss.v059.i05.

36. Aronoff, J.E., et al. Evidence for an energy conservation model of inflammaging and immunosenescence in the US Health and Retirement Study and UK Biobank. The Journals of Gerontology, Series A: Biological Sciences and Medical Sciences. 2026;10.1093/gerona/glag119.

37. Aronoff, J.E. and B.C. Trumble. An evolutionary medicine and life history perspective on aging and disease: Trade-offs, hyperfunction, and mismatch. Evol Med Public Health. 2025;13:111–124. 10.1093/emph/eoaf010.

38. Sorrenti, V., et al. Immunomodulatory and Antiaging Mechanisms of Resveratrol, Rapamycin, and Metformin: Focus on mTOR and AMPK Signaling Networks. Pharmaceuticals (Basel). 2022;15:912. 10.3390/ph15080912.

39. Wang, R., B. Sunchu, and V.I. Perez. Rapamycin and the inhibition of the secretory phenotype. Exp Gerontol. 2017;94:89–92. 10.1016/j.exger.2017.01.026.

40. Lin, H., et al. The Role and Mechanism of Metformin in Inflammatory Diseases. J Inflamm Res. 2023;16:5545–5564. 10.2147/JIR.S436147.

41. Mehdi, S.F., et al. Glucagon-like peptide-1: a multi-faceted anti-inflammatory agent. Front Immunol. 2023;14:1148209. 10.3389/fimmu.2023.1148209.

42. Cohen, A.A. and M. Picard. Potential Risks of Blocking GDF15-Based Brain Energy Sensing. J Am Geriatr Soc. 2026;10.1111/jgs.70399.

43. Muehlenbein, M.P., et al. Lower testosterone levels are associated with higher risk of death in men. Evol Med Public Health. 2023;11:30–40. 10.1093/emph/eoac044.

44. Kraft, T.S., et al. Nutrition transition in 2 lowland Bolivian subsistence populations. Am J Clin Nutr. 2018;108:1183–1195. 10.1093/ajcn/nqy250.

45. Gurven, M., et al. The Tsimane Health and Life History Project: Integrating anthropology and biomedicine. Evol Anthropol. 2017;26:54–73. 10.1002/evan.21515.

46. Kaplan, H., et al. Coronary atherosclerosis in indigenous South American Tsimane: a cross-sectional cohort study. Lancet. 2017;389:1730–1739. 10.1016/S0140-6736(17)30752-3.

47. Franck, M., et al. Nonuniversality of inflammaging across human populations. Nat Aging. 2025;5:1471–1480. 10.1038/s43587-025-00888-0.

48. Aronoff, J.E., et al. Inflammaging is minimal among forager-horticulturalists in the Bolivian Amazon. Proceedings of the Royal Society B: Biological Sciences. 2025;292:10.1098/rspb.2025.1111.

49. Franck, M., et al. Inflamm-aging as a diverse and context-dependent process: from species and population differences to individual trajectories. Ageing Research Reviews. 2025;102880.

50. Trumble, B.C., et al. Challenging the inevitability of prostate enlargement: low levels of benign prostatic hyperplasia among Tsimane Forager-Horticulturalists. Journals of Gerontology Series A: Biomedical Sciences and Medical Sciences. 2015;70:1262–1268. 10.1093/gerona/glv051.

51. Trumble, B.C., et al. Testosterone is positively associated with coronary artery calcium in a low cardiovascular disease risk population. Evolution, Medicine, and Public Health. 2023;11:472–484. 10.1093/emph/eoad039.

